# Differentiation of rhizosphere fungal assemblages by host ploidy level in mixed-ploidy *Larrea tridentata* populations

**DOI:** 10.1101/2022.11.04.515195

**Authors:** Benjamin Gerstner, Michael Mann, Robert G. Laport, Kenneth D. Whitney

**Affiliations:** Department of Biology, University of New Mexico, Albuquerque, NM 87131, USA; Department of Biology, The College of Idaho, Caldwell, ID 83605, USA

**Keywords:** polyploidy, minority cytotype exclusion, chromosome doubling, rhizosphere communities, *Larrea tridentata*, niche differentiation, indicator species

## Abstract

Polyploidy—whole genome duplication—is common in plants. Studies over the last several decades have documented numerous mixed-ploidy populations. Whether arising via recurrent whole genome duplication events within a population, or from secondary contact, the persistence of mixed populations depends on the ability of the minority cytotype to overcome the negative frequency dependent effects of outcrossing with other ploidies, known as Minority Cytotype Exclusion. One mechanism of overcoming Minority Cytotype Exclusion is microbially-mediated niche differentiation (MMND), wherein cytotypes occupy different niches via interactions with different sets of microbes. Inherently cryptic, MMND is underexplored in polyploid plant populations. Here, we search for evidence of MMND in creosotebush (*Larrea tridentata*), a dominant desert shrub of the southwestern U.S. and northern Mexico. We sequenced fungi from rhizosphere soils of diploid, autotetraploid, and autohexaploid plants growing in two naturally-occurring mixed-cytotype populations. Within populations, we found substantial fungal assemblage overlap across host plant cytotypes. However, using indicator species analysis, we identified some fungi that are differentiated by host plant cytotype, satisfying a precondition for MMND. Future study is needed to determine the degree of niche differentiation conferred, if any, and whether the identified fungi play a role in the long-term persistence of multiple cytotypes within populations.

## Introduction

Angiosperms have a long history of polyploidization (Landis et al. 2018), the origination and maintenance of more than two complete chromosome sets within an organism. Within a single species, polyploid complexes can form via recurrent polyploidization events within a population resulting in multiple cytotypes (e.g., diploid, tetraploid, etc.) occurring at the same location. Over the last several decades, there has been a renewed interest in understanding the population-level processes driving cytotype co-occurrence and patterns of biodiversity (Coyne and Orr 2004; Ramsey and Ramsey 2014; Segraves and Anneberg 2016; Laport and Ng 2017). Theoretical predictions indicate mixed-cytotype populations should be short-lived due to Minority Cytotype Exclusion (MCE). MCE posits that because a newly arising cytotype is both rare compared to its progenitor cytotype and reproductively incompatible with it, the newly arising cytotype will have low fitness. Therefore, the minority cytotype will typically be excluded – a form of frequency-dependent selection (Levin 1975). However, the presence of numerous polyploid lineages (Wood et al. 2009) suggests that MCE is far from an inevitable outcome for many polyploid complexes.

Theory suggests that niche differentiation is one mechanism by which minority cytotypes may overcome MCE (Fowler and Levin 2016). Niche differentiation among cytotypes has been documented for many species and is linked to both abiotic niche axes (e.g., substrate, elevation, temperature, moisture; Laport et al. 2013; López-Jurado, Mateos-Naranjo, and Balao 2019; Wan et al. 2019; Decanter et al. 2020) and biotic niche axes (e.g., herbivores, pollinators; Münzbergová, Skuhrovec, and Maršík 2015; Laport et al. 2016; Muñoz-Pajares et al. 2018; Čertner et al. 2019; O’Connor, Laport, and Whiteman 2019; Laport, Minckley, and Pilson 2021).

Even slight differences in traits between diploids and polyploids may facilitate successful escape from MCE (B. Husband 2000) by easing direct ecological competition and promoting assortative mating. Such differences may arise upon formation of new cytotypes, or through a period of post-polyploidization isolation and adaptation to novel ecological conditions prior to secondary contact. The diversity of phenotypic differences that have been documented between diploids and polyploids (Levin 1983; Segraves and Thompson 1999; B. C. Husband et al. 2008; Maherali, Walden, and Husband 2009; López-Jurado, Mateos-Naranjo, and Balao 2019) may help explain why so many present-day plant communities contain polyploids co-occurring with their close diploid relatives (Gaynor et al. 2018).

Microbially-mediated niche differentiation (MMND) represents a possible cryptic and underexplored mechanism of niche differentiation for polyploid complexes. In MMND, microbes can help plants acquire nutrients and thus expand or shift their niche dimensions. For example, derived cytotypes (e.g., tetraploids, hexaploids, etc.) can have different quantities of root exudates (Wu et al. 2019), which may allow them to recruit distinctive microbial communities (Segraves 2017). Prior research on mixed-cytotype populations of orchids in the *Gymnadenia conopsea* group have found cytotype-specific root-associated fungal assemblages (Těšitelová et al. 2013). This observation held for both field-collected adults and seedlings and was most pronounced within a site with closely sympatric adults (within 1m^2^). In close proximity, diploid *G. conopsea* shared only one occurrence of the same fungal OTU with tetraploid *G. conopsea* (Těšitelová et al. 2013). This compositional difference in root-associated fungal assemblages may contribute to different niche occupation (*sensu* MacArthur 1958) and an easing of MCE that allows coexistence of these orchid cytotypes. In contrast, observations from *Aster amellus* and *Centaurea stoebe* indicate no significant differences in arbuscular mycorrhizal fungi between diploids and tetraploids, suggesting root mycorrhizal associations may not be strong contributors to niche differentiation in all polyploid species(Sudova et al. 2014; Sudová et al. 2018). Thus, it remains unclear if MMND is common among taxa comprising multiple cytotypes.

The North American creosote bush [*Larrea tridentata* (DC.) Coville; Zygophyllaceae] is an autopolyploid complex distributed across the southwestern U.S. and northern Mexico (Mabry, Hunziker, and Difeo, Jr. 1977). The complex comprises three distinct cytotypes (diploid, 2*n* = 2*x* = 26; tetraploid, 2*n* = 4*x* = 52; and hexaploid, 2*n* = 6*x* = 78) with distributions approximately corresponding to the three warm deserts of North America (Chihuahuan Desert, Sonoran Desert and Mojave Desert, respectively) in which they are dominant shrubs (Mabry, Hunziker, and Difeo 1977). Prior work has mapped the cytogeography of the complex, identifying multiple natural contact zones and broad overlap of the 4*x* and 6*x* cytotypes in the Sonoran Desert (Hunter et al. 2001; Yang 1970; Laport, Minckley, and Ramsey 2012; Laport and Ramsey 2015). The current distributions with narrow areas of cytotype contact likely represent secondary contact after complex biogeographic histories involving migration and adaptation during glacial and post glacial periods (Hunter et al. 2001; Laport, Minckley, and Ramsey 2016), though polyploid cytotypes may be recurrently formed and could represent instances of primary contact (Laport, Minckley, and Ramsey 2016). Although the cytotypes appear to be at least partially ecologically differentiated along several niche axes (e.g., climatic, vegetation communities, herbivore specificity, pollinator visitation; Laport et al. 2013; Laport, Minckley, and Ramsey 2016; O’Connor, Laport, and Whiteman 2019; Laport, Minckley, and Pilson 2021) it remains unclear whether such differences are sufficient to maintain sympatry or whether MMND may interact with other niche differences to ease MCE at natural areas of contact.

Here, we investigate whether MMND might play a role in easing MCE and facilitating cytotype co-occurrence at naturally occurring 2*x*-4*x* and 4*x*-6*x* mixed-cytotype populations. We hypothesize that in mixed-cytotype populations cytotypes have largely dissimilar rhizosphere fungal assemblages and exhibit host-cytotype rhizosphere fungal specialization consistent with MMND.

## Materials and Methods

### Study system

*Larrea tridentata* is a long-lived perennial evergreen shrub that reproduces via seed, but may also propagate clonally (Mabry, Hunziker, and Difeo, Jr. 1977). The three cytotypes have relatively well-defined distributions, likely maintained by abiotic environmental variation, but also potentially determined by pollinator-mediated assortative mating or galling midge interactions at cytotype contact zones (Laport, Minckley, and Pilson 2021; O’Connor, Laport, and Whiteman 2019). In mixed-cytotype populations, 4*x L. tridentata* tend to be found in denser vegetation associations than 2*x* or 6*x* plants, which tend to be found at higher elevations, in more species-rich communities and on coarser soils (Laport, Minckley, and Ramsey 2016). Tetraploids tend to flower earlier and produce more flowers than 2*x* or 6*x* plants (Laport, Minckley, and Ramsey 2016). The size of morphological structures tends to increase with ploidy (e.g., larger diameter pollen grains, longer stamens and pistils, and longer, wider petals and leaves) though 4*x* plants tend to be taller than either 2*x* or 6*x* plants (Laport, Minckley, and Ramsey 2016). In mixed-cytotype 2*x*-4*x* populations, 4*x* plants have a significantly higher rate of bee visitation, but 2*x* pollen is over-represented on native bees, which may contribute to assortative mating and the maintenance of cytotype coexistence (Laport, Minckley, and Pilson 2021). The distributions of specialist herbivore species have also been documented to be concordant with 2*x* and 4*x L. tridentata*, potentially resulting in cytotype-specific fitness differences that may also enable narrow zones of cytotype co-occurrence (O’Connor, Laport, and Whiteman 2019).

### Field collections and sample preparation

Prior research leveraging flow-cytometric analyses to infer DNA content has identified multiple mixed-cytotype *L. tridentata* populations comprising permanently marked plants (Laport, Minckley, and Ramsey 2012, Laport and Ramsey 2015). In the current work, we sampled rhizosphere soils under plants of known ploidy from one 2*x*-4*x* population (San Pedro 3; 32.60°, -110.54°) and two 4*x*-6*x* populations (Algodones N4; 33.00°, -115.07° and Algodones S3; 32.81°, -114.87°; Laport and Ramsey 2015; see Fig. 1). Because the mixed-cytotype sites are asymmetrically mixed, we combined the two 4*x*-6*x* populations in our analyses (and hereafter refer to them as a single population -Algodones) to ensure large enough rhizosphere sample sizes for each co-occurring ploidy. In April 2021, rhizosphere soils were collected from multiple holes dug inward near the shrub base until fine roots were observed (minimum of 30cm deep) and pooled (to obtain a minimum of ∼10mL of soil). In total 10 diploid, 27 tetraploid and 17 hexaploid rhizosphere samples were collected.

**Figure 1:**
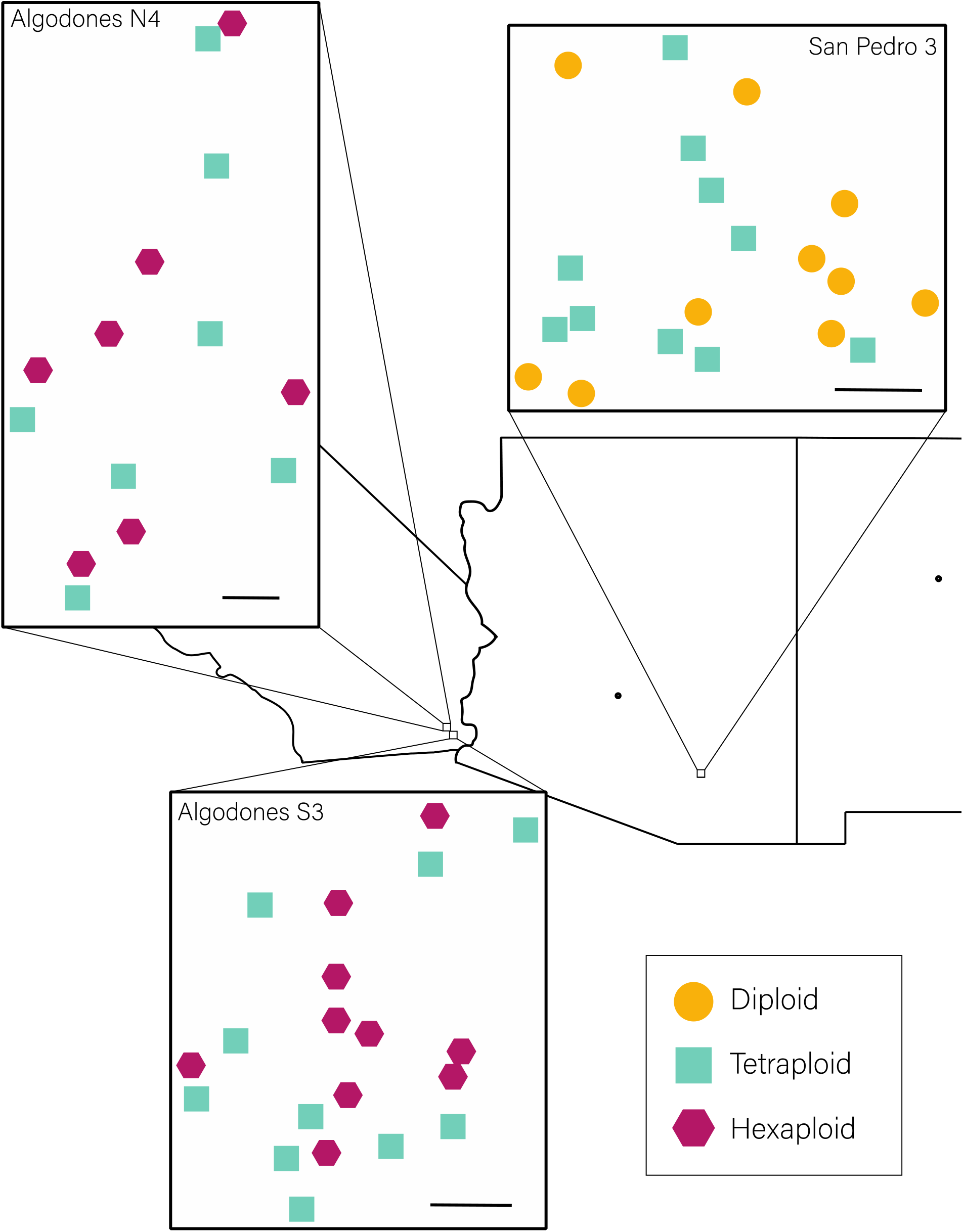
Localities for 54 rhizosphere collections of 2*x* (circles), 4*x* (squares) and 6*x* (hexagons) *Larrea tridentata* from southeastern California and southern Arizona. A 100 m scale is represented for each collection site (black bar).

Soils were stored on ice and refrigerated for up to 72 h prior to performing DNA extractions. DNA extractions were performed in randomized batches with Qiagen PowerSoil kits (Qiagen catalog number 47014) and completed within a two-day period at the University of New Mexico. Extraction quality and DNA yield was assessed using a Qubit 3.0 Fluorometer (ThermoFisher Scientific).

### Molecular and bioinformatic work

Fungal DNA in each soil extract was amplified with fungal-specific primers (ITS3-FUN and ITS4-FUN) spanning the ITS2 region (Taylor et al. 2016). Each reaction used the following PCR machine conditions using Phusion polymerase: First, an initial denaturation at 98°C for 30 seconds, followed by 27 cycles of annealing at 58°C for 10s, extension for 60°C for 4 minutes, and then concluded with a final extension at 60°C for 20 minutes. A second PCR was conducted to add the nextera adaptors following the same conditions but for 7 cycles. Each sample was pooled at equal concentration and sequenced on 2 × 300 cycle Illumina MiSeq runs. The forward and reverse reads were merged using USEARCH9 (Edgar 2010). The primer regions were removed using Cutadapt 3.5 (Martin 2011). The sequences were then filtered to include less than one expected error, and then clustered into OTUs at 97% similarity with UPARSE (Edgar 2013). The taxonomy for each OTU was determined by running SINTAX (Edgar 2016) from USEARCH and using Utax 8.2 (Abarenkov et al. 2021) as the reference dataset. The OTU table was created by mapping reads to OTUs using the *usearch_global* function in USEARCH. The non-fungal OTUs were removed from the OTU table and then rarefied to 1000 reads in R 4.1 (R Core Team 2022) using *EcoUtils* package (Salazar 2022).

### Statistical analyses

We tested for differences in fungal assemblage composition using Bray-Curtis dissimilarities and a PERMANOVA model (composition ∼ ploidy + population + ploidy*population) via the *adonis2* function in the R 4.1 package *vegan* (Table S1, Oksanen et al. 2020). Ploidy was coded as ‘ancestral’ and ‘derived’, permitting all data from both mixed sites to be analyzed together.

For example, rhizosphere samples from 4*x* plants were coded as ‘derived’ at the 2*x*-4*x* site but as ‘ancestral’ at the 4*x*-6*x* site. ‘Ancestral’ indicates a cytotype that gave rise to a ‘derived’ cytotype at some time in the past, but we do not assume that the ‘derived’ cytotype necessarily formed in the studied populations. The fixed factors were Ploidy and Population, the test was to measure marginal effects, and the number of permutations was set to 10,000. To visualize potentially discrete fungal assemblages, we utilized non-metric multidimensional scaling (NMDS) via the *amp_ordinate* function from *ampvis2* (Andersen et al. 2018) and visualized using ggplot2 (Wickham 2016).

Indicator species analysis was performed on the OTU dataset (coded as presence/absence) using the *indicspecies* package (Cáceres and Legendre 2009) in R, with each plant designated as a ‘site’ and plant cytotype as the ‘type’ of site. This analysis resulted in indicator value indices, which can be parsed to two conditional probabilities, specificity and fidelity (Dufrêne and Legendre 1997; Cáceres and Legendre 2009). Specificity is the probability that a plant belongs to a given cytotype, given an association with a particular fungal OTU. Fidelity is the probability of finding a given fungal OTU in association with a particular cytotype. Specificity and fidelity values aid in determining that specialization is present, and whether specialization is genuine and not simply because fungal OTUs are rare or present in the dataset due to sequencing artifacts.

Theory further suggests that while some individual species may not be indicators alone, combinations of a few species (e.g., pairs, triplets, etc.) may be indicators of ecological/niche differences (De Cáceres et al. 2012; see Bouffaud et al. 2017 for evidence in arbuscular mycorrhizal fungi). We thus tested for significant indicator OTUs and OTU combinations (up to four) that occurred in rhizosphere samples associated with the three cytotypes of *L. tridentata* using the *indicators* and *pruneindicators* functions in the *indicspecies* R package. Confidence intervals were calculated from 10,000 bootstrap replicates. Putative functional assignments for indicator OTUs were made by querying the FungalTraits V1.2 database (Põlme et al. 2020). All R code used the *tidyverse* package (Wickham et al. 2019).

## Results

In total we detected 2217 OTUs (1569 in San Pedro 3, 654 in Algodones N4, 944 in Algodones S3; 1057, 1667, 868 in 2*x*, 4*x* and 6*x* rhizospheres, respectively). In mixed-cytotype populations, rhizosphere fungal assemblages exhibited considerable overlap in functional group assignments and did not significantly differ between cytotypes (Figs. 2A, 2B; Table S1). Rather than cytotype, population identity was the strongest explanatory variable for rhizosphere fungal assemblage composition (Table S1, Fig. S1). These results were robust across different read-number filtering choices (Table S1). Comparisons were made of normalized read counts for the top 75 OTUs and none significantly differed between pairs of cytotypes (Figs. 3A, 3B; Fig. S2).

**Figure 2:**
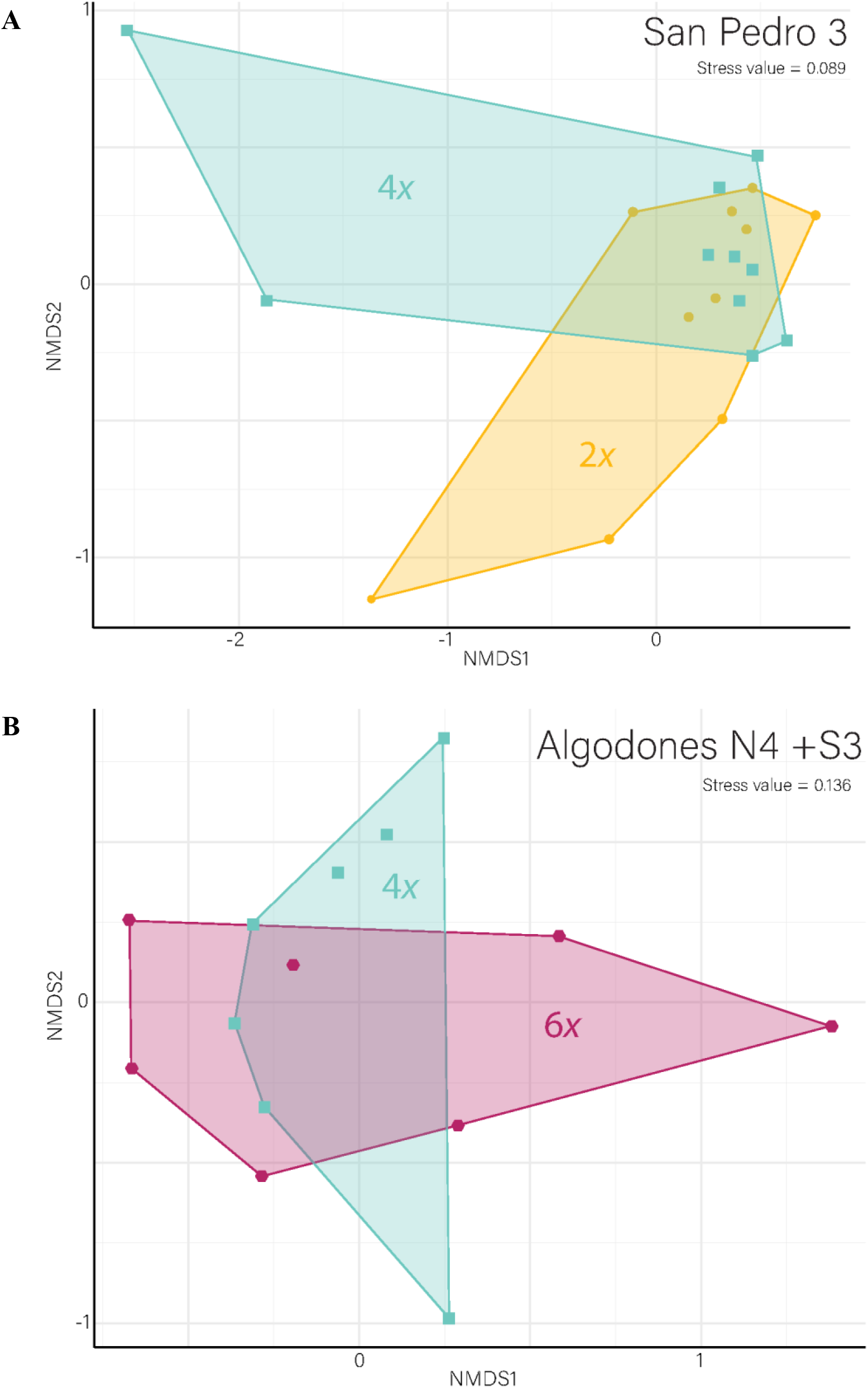
NMDS ordination for fungal assemblage composition as a function of ploidy for 2*x*-4*x* (**A**, 2*x* = yellow, 4*x* = blue) and 4*x*-6*x* (**B**, 4*x* = blue, 6*x* = red) *Larrea tridentata* populations. Each point represents the fungal assemblage composition for a single plant rhizosphere. Rhizosphere fungal assemblages exhibit some overlap between ploidies for both populations, but the fungal assemblages are not concordant.

**Figure 3:**
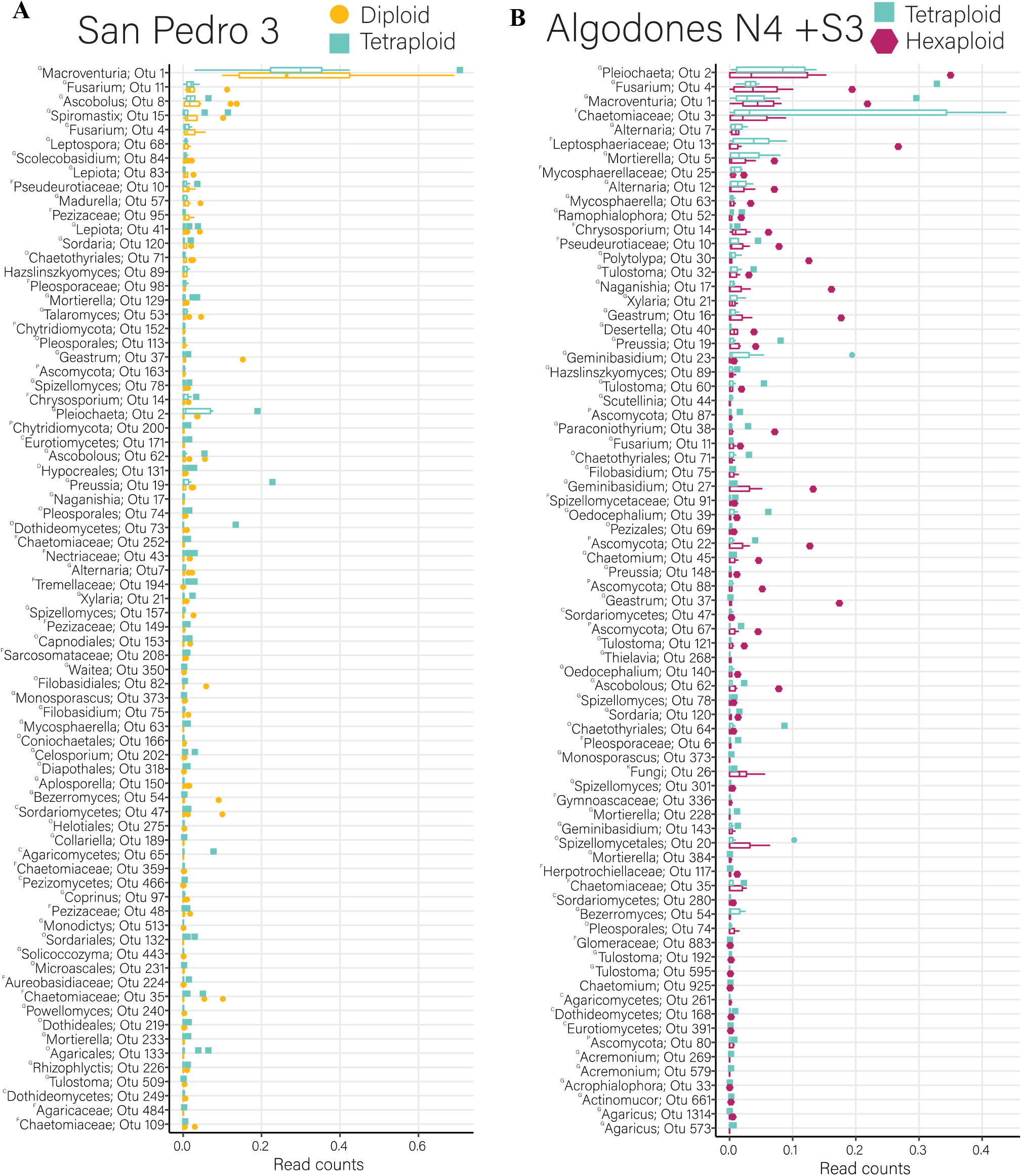
Top 75 fungal OTUs, by normalized read counts, for 2*x*-4*x* (**A**, 2*x* = yellow, 4*x* = blue) and 4*x*-6*x* (**B**, 4*x* = blue, 6*x* = red) populations. Boxplots show the median and interquartile range with outliers shown as dots. Taxa are ordered by highest median normalized read count to lowest. The lowest degree of biological classification is named for each OTU, and the leading superscripts identify the classification level, (^K^ – Kingdom, ^P^ – Phylum, ^C^ – Class, ^O^ – Order, ^F^ – Family, ^G^ – Genus). None of the normalized read counts are significantly different between ploidies.

Indicator species analyses revealed differences between cytotypes in rarer fungal taxa that were not identified in the assemblage-level analyses. Despite the degree of overlap in fungal OTUs between co-occurring cytotypes, our indicator species analyses suggested the cytotypes in both mixed-cytotype populations had at least some unique fungal associations (Tables 1 & 2). Four indicator OTUs were identified in the 2*x*-4*x* population (San Pedro; two indicating 2*x*, and two indicating 4*x*), while 12 indicator OTUs were identified in the combined 4*x*-6*x* populations (Algodones; eight indicating 4*x*, and four indicating 6*x*). We further identified combinations of fungal OTUs with strong support as indicator combinations for *L. tridentata* cytotypes (Tables 3 & 4). Four indicator combinations were identified in the San Pedro population (two indicating 2*x*, and two indicating 4*x*), while three were identified in the Algodones population (one indicating 4*x*, and two indicating 6*x*). We were unable to make specific functional assignments from the FungalTraits database for some of the indicator OTUs on 2*x L. tridentata*, but those that could be assigned were mostly wood or soil saprotrophs (Tables 1 & 3). Functional assignments for 4*x* plants were mostly soil or dung saprotrophs (Tables 1, 2, 3, 4), and those for 6*x* plants were mostly litter saprotrophs (Tables 2 & 4).

**Table 1:** Indicator OTUs, by ploidy, for the San Pedro 2*x*-4*x Larrea tridentata* population (San Pedro 3) based on presence/absence data. Ploidy is either diploid (2*x*) or tetraploid (4*x*) and is the cytotype for which the OTU is an indicator. Specificity is the probability that the plant belongs to that cytotype, given that the OTU has been found there. Fidelity is the probability of finding that OTU on that cytotype. Values in parentheses are 95% confidence intervals based on 10,000 bootstrap replicates. Bolded levels of classification came from searches on NCBI BLAST, all others are from SINTAX/USEARCH results. Primary/Secondary Lifestyle are from FungalTraits 1.2V.

| *Ploidy* | *Specificity* | *Fidelity* | *p-value* | *OTU* | Phylum;<br>Class;<br>Order | Family;<br>*Genus species*<br>Primary/Secondary Lifestyle |
| --- | --- | --- | --- | --- | --- | --- |
| 2x | 1<br>(1-1) | 0.5<br>(0.182-0.818) | 0.032 | 631 | Chytridiomycota;<br>Spizellomycetes;<br>Spizellomycetales | -;<br>-<br>Unable to determine |
| 2x | 1<br>(1-1) | 0.5<br>(0.182-0.818) | 0.033 | 1988 | **Ascomycota;**<br>**Leotiomycetes;**<br>**Helotiales** | -;<br>-<br>Unable to determine |
| 4x | 1<br>(1-1) | 0.6<br>(0.273-0.9) | 0.010 | 1777 | Ascomycota;<br>Dothideomycetes;<br>Pleosporales | Pleosporaceae;<br>*Curvularia sp.*<br>Plant pathogen/Litter saprotroph |
| 4x | 1<br>(1-1) | 0.5<br>(0.182-0.818) | 0.033 | 52 | Ascomycota;<br>**Sordariomycetes;**<br>**Sordariales** | **Lasiosphaeriaceae;**<br>***Ramophialophora sp.***<br>Soil saprotroph |

**Table 2:** Indicator OTUs, by ploidy, for the Algodones 4*x*-6*x Larrea tridentata* population (Algodones N4 + S3) based on presence/absence data. Ploidy is either tetraploid (4*x*) or hexaploid (6*x*) and is the cytotype for which the OTU is an indicator. Specificity is the probability that the plant belongs to that cytotype, given that the OTU has been found there. Fidelity is the probability of finding that OTU on that cytotype. Values in parentheses are 95% confidence intervals based on 10,000 bootstrap replicates. Bolded levels of classification came from searches on NCBI BLAST, all others are from SINTAX/USEARCH results. Primary/Secondary Lifestyle are from FungalTraits 1.2V.

| *Ploidy* | *Specificity* | *Fidelity* | *p-value* | *OTU* | Phylum;<br>Class;<br>Order; | Family;<br>*Genus species*<br>Primary/Secondary Lifestyle |
| --- | --- | --- | --- | --- | --- | --- |
| 4x | 0.857<br>(0.5-1) | 0.462<br>(0.188-0.750) | 0.018 | 1030 | Basidiomycota;<br>Cystobasidiomycetes;<br>Cystobasidiales | Cystobasidiaceae;<br>*Cystobasidium pallidum*<br>Mycoparasite |
| 4x | 1<br>(1-1) | 0.385<br>(0.125-0.667) | 0.008 | 393 | Ascomycota;<br>Pezizomycetes;<br>Pezizales | Pezizaceae;<br>*Iodophanus sp.*<br>Soil/Dung saprotroph |
| 4x | 1<br>(1-1) | 0.308<br>(0.077-0.583) | 0.035 | 542 | **Chytridiomycota;**<br>**Chytridiomycetes;**<br>- | -;<br>-<br>Unable to determine |
| 4x | 1<br>(1-1) | 0.308<br>(0.071-0.571) | 0.035 | 394 | Ascomycota;<br>Dothideomycetes;<br>Pleosporales | Sporormiaceae;<br>***Preussia sp.***<br>Dung saprotroph |
| 4x | 1<br>(1-1) | 0.308<br>(0.077-0.571) | 0.031 | 486 | Ascomycota;<br>Sordariomycetes;<br>Sordariales | Podosporaceae;<br>***Podospora sp.***<br>Dung saprotroph |
| 4x | 1<br>(1-1) | 0.308<br>(0.071-0.571) | 0.034 | 818 | Ascomycota;<br>Dothideomycetes;<br>Pleosporales | Sporormiaceae;<br>*Preussia sp.*<br>Dung saprotroph |
| 4x | 1<br>(1-1) | 0.308<br>(0.071-0.583) | 0.029 | 2567 | Ascomycota;<br>Dothideomycetes;<br>Pleosporales | Didymosphaeriaceae;<br>*Deniquelata sp.*<br>Plant pathogen/Soil saprotroph |
| 4x | 1<br>(1-1) | 0.308<br>(0.077-0.571) | 0.032 | 1970 | **Basidiomycota;**<br>**Cystobasidiomycetes;**<br>**Cystobasidiales** | **Cystobasidiaceae;**<br>***Cystobasidium sp.***<br>Mycoparasite |
| 6x | 0.625<br>(0.429-0.815) | 1<br>(1-1) | 0.01 | 45 | Ascomycota;<br>Sordariomycetes;<br>Sordariales | Chaetomiaceae;<br>*Chaetomium sp.*<br>Litter/Foliar saprotroph |
| 6x | 0.786<br>(0.538-1) | 0.733<br>(0.5-0.938) | 0.006 | 201 | Ascomycota;<br>Sordariomycetes;<br>Hypocreales; | Cordycipitaceae;<br>-<br>Unable to determine |
| 6x | 0.818<br>(0.556-1) | 0.6<br>(0.333-0.846) | 0.011 | 1566 | Ascomycota;<br>Sordariomycetes;<br>- | -;<br>-<br>Unable to determine |
| 6x | 1<br>(1-1) | 0.333<br>(0.1-0.583) | 0.043 | 2201 | Ascomycota;<br>Pezizomycetes;<br>Pezizales | Ascobolaceae;<br>*Ascobolus sp.*<br>Litter/Dung saprotroph |

**Table 3:** Indicator OTU combinations, by ploidy, for the San Pedro 2*x*-4*x Larrea tridentata* population (San Pedro 3) based on presence/absence data. Ploidy is diploid (2*x*) or tetraploid (4*x*) and is the cytotype for which the OTU combination is an indicator. Specificity is the probability that the plant belongs to that cytotype, given that the OTU combination has been found there. Fidelity is the probability of finding that OTU combination on that cytotype. Values in parentheses are 95% confidence intervals based on 10,000 bootstrap replicates. Bolded levels of classification came from searches on NCBI BLAST, all others are from SINTAX/USEARCH results. Primary/Secondary Lifestyle are from FungalTraits 1.2V.

| *Ploidy* | *Specificity* | *Fidelity* | *p-value* | *OTU*<br>*Combination* | Phylum;<br>Class;<br>Order | Family;<br>*Genus species*<br>Primary/Secondary Lifestyle |
| --- | --- | --- | --- | --- | --- | --- |
| 2x | 0.727<br>(0.444-1) | 0.8<br>(0.5-1) | 0.035 | 13 | Ascomycota;<br>Dothideomycetes;<br>Pleosporales | Leptosphaeriaceae;<br>***Leptosphaeria sp.***<br>Plant pathogen/Wood saprotroph |
|  |  |  |  | 45 | Ascomycota;<br>Sordariomycetes;<br>Sordariales | Chaetomiaceae;<br>*Chaetomium sp.*<br>Litter/Foliar saprotroph |
|  |  |  |  | 2464 | Basidiomycota;<br>Tremellomycetes;<br>**Filobasidiales** | **Filobasidiaceae;**<br>***Naganishia sp.***<br>Unspecified saprotroph |
| 2x | 0.727<br>(0.444-1) | 0.8<br>(0.5-1) | 0.035 | 25 | Ascomycota;<br>Dothideomycetes;<br>Capnodiales | Mycosphaerellaceae;<br>-<br>Unable to determine |
|  |  |  |  | 38 | **Ascomycota;**<br>**Dothideomycetes;**<br>**Pleosporales** | **Didymellaceae;**<br>***Paraconiothyrium sp.***<br>Wood saprotroph |
|  |  |  |  | 53 | Ascomycota;<br>Eurotiomycetes;<br>Eurotiales | Trichocomaceae;<br>*Talaromyces sp.*<br>Unspecified saprotroph |
|  |  |  |  | 64 | Ascomycota;<br>Eurotiomycetes;<br>Chaetothyriales | **Trichocomaceae;**<br>***Knufia sp.***<br>Soil saprotroph |
| 4x | 0.778<br>(0.5-1) | 0.7<br>(0.4-1) | 0.020 | 2 | Ascomycota;<br>Dothideomycetes;<br>Pleosporales | Didymellaceae;<br>***Pleiochaeta setosa***<br>Plant pathogen/Wood saprotroph |
|  |  |  |  | 19 | Ascomycota;<br>Dothideomycetes;<br>Pleosporales | Sporormiaceae;<br>*Preussia sp.*<br>Dung saprotroph |
|  |  |  |  | 151 | Ascomycota;<br>Pezizomycetes;<br>Pezizales | **Pyronemataceae;**<br>-<br>Unable to determine |
| 4x | 0.778<br>(0.455-1) | 0.7<br>(0.375-1) | 0.045 | 37 | Basidiomycota;<br>Agaricomycetes;<br>Geastrales | Geastraceae;<br>*Geastrum sp.*<br>Litter saprotroph |
|  |  |  |  | 103 | Ascomycota;<br>Dothideomycetes;<br>**Pleosporales** | -;<br>-<br>Unable to determine |
|  |  |  |  | 169 | **Ascomycota;**<br>**Dothideomycetes;**<br>- | -;<br>-<br>Unable to determine |

**Table 4:**
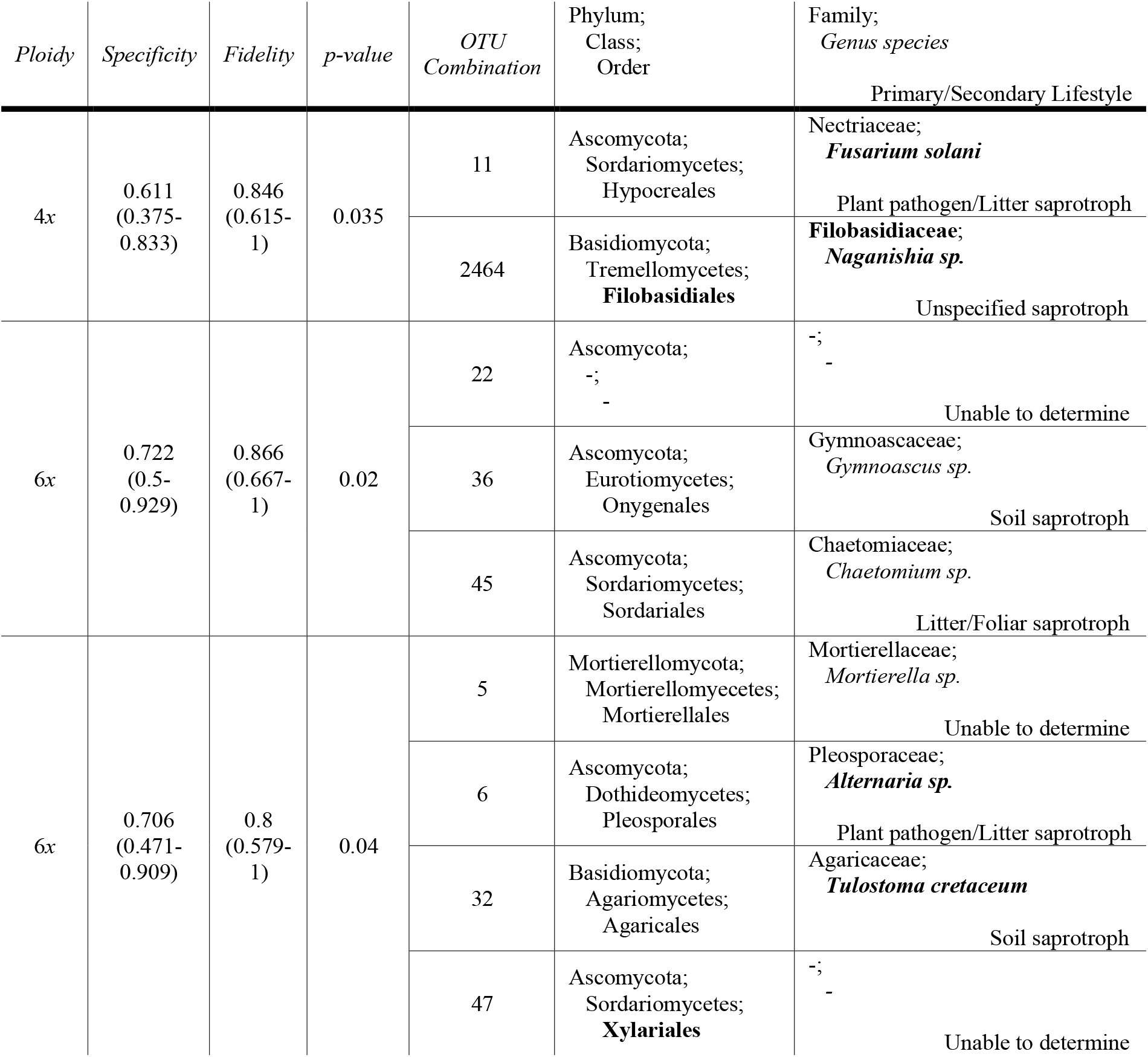
Indicator OTU combinations, by ploidy, for the Algodones 4*x*-6*x Larrea tridentata* population (Algodones N4 + S3) based on presence/absence data. Ploidy is either tetraploid (4*x*) or hexaploid (6*x*) and is the cytotype for which the OTU combination is an indicator. Specificity is the probability that the plant belongs to that cytotype, given that the OTU combination has been found there. Fidelity is the probability of finding that OTU combination on that cytotype. Values in parentheses are 95% confidence intervals based on 10,000 bootstrap replicates. Bolded levels of classification came from searches on NCBI BLAST, all others are from SINTAX/USEARCH results. Primary/Secondary Lifestyle are from FungalTraits 1.2V.

## Discussion

Rare cytotypes in mixed-cytotype populations face extinction due to minority cytotype exclusion (MCE). However, mixed-cytotype populations have increasingly been documented over the last several decades with the advent of high-throughput flow cytometry, suggesting many newly-arisen polyploids overcome MCE. One mechanism by which MCE may be overcome, facilitating co-occurrence among cytotypes, is niche differentiation. MMND is a hypothesized cryptic means of niche differentiation for polyploids through the differential associations of microbes recruited to plants of different ploidy. Here we found that naturally co-occurring diploid, tetraploid, and hexaploid cytotypes of *L. tridentata* exhibited broad overlap in rhizosphere fungal associates. Yet, we also found support for a precondition of microbially mediated niche differentiation in both 2*x*-4*x* and 4*x*-6*x* mixed-cytotype populations as diploids, tetraploids, and hexaploids each had some unique fungal associates.

### Understanding rhizosphere fungal assemblage overlap and host cytotype specialization

We hypothesized that cytotypes from mixed-cytotype populations would have differentiated rhizosphere fungal assemblages but found that total rhizosphere fungal assemblages were similar for co-occurring cytotypes. Ordination of the sampled fungal OTUs suggested very little differentiation among fungal assemblages on co-occurring diploids and tetraploids (Fig. 2A) or co-occurring tetraploids and hexaploids (Fig. 2B), and there was no support for assemblage differences in our PERMANOVA (Table S1). Thus, plants within a population had very similar total rhizosphere fungal assemblage regardless of cytotype.

We further hypothesized that there would be rhizosphere fungal specialization to host cytotype. We found that cytotypes from both mixed-cytotype populations were associated with distinctive indicator OTUs, suggesting niche differentiation that is consistent with MMND (Tables 1-4). Though cryptic, such niche differentiation could help facilitate cytotype co-occurrence by easing MCE.

At first, observing no major differences between rhizosphere fungal assemblages among cytotypes seems incongruent with finding strong support for host cytotype specialization. One possible explanation follows from the fact that the same data are analyzed differently by PERMANOVA and indicator species analyses. PERMANOVA results are driven by abundant species (as measured by normalized read counts), whereas indicator species analysis is driven by unique species occurrences. We surmise there is a ‘core’ rhizosphere fungal assemblage of abundant taxa associated with *L. tridentata* regardless of ploidy; in addition, there appears to be a group of rarer fungal taxa that constitute the distinctive combinations of indicator OTUs that differ by ploidy.

When using non-identity metrics, prior studies failed to find evidence of cytotype-specific root-associated fungal differences between ploidies. In *Aster amellus* and *Centaurea stoebe* (Sudová et al. 2014; Sudová et al. 2018) there was no significant difference between ploidies in arbuscular mycorrhizal root colonization or extraradical mycelium length. Regardless of finding no cytotype specific differences, without using identity metrics it is not possible to say whether the precondition for MMND is met or not in these two systems. A different study that used fungal identities has documented differences in root-fungal associations between co-occurring diploid and tetraploid *Gymnadenia conopsea* orchids (Těšitelová et al. 2013). They found cytotype-specific OTUs with the highest relative abundances predominately belonged to the Tulasnellaceae, which commonly form endomycorrhizal associations with orchids. Five of these cytotype-specific OTUs showed evidence of being distinct from *Tulasnella* reference species in GenBank and may be most closely related to a known wood saprotroph, *Gleotulasnella cystidiophora*. These findings support the precondition of cytotype-specific differences for MMND.

Our work differs from prior studies (Sudova 2014, Sudova 2018) in our use of fungal identity-based measures and (Tesitelova 2013) in our use of rhizosphere soils. Sequencing rhizosphere soils where fine root pieces are present means we are identifying a more complete root-associated fungal assemblage (both endo-and ectomycorrhizal associations), whereas prior studies have only evaluated endomycorrhizal associations (Teitelova 2013, Sudova 2014, Sudova 2018).

### Potential functional differences derived from host-specialized OTUs

Attributing function to host-specialized OTUs allows us to hypothesize about how rhizosphere microbial interactions might contribute to niche differentiation (MMND) between *L. tridentata* cytotypes. Primary and secondary lifestyle assignment of identified fungal OTUs come from the FungalTraits V1.2 database (Tables 1-4) and are specific to the OTU genus. These assignments are useful to consider as there is a wide variety of possible interactions between the rhizosphere fungi and their hosts. For example, the many indicator OTUs for the tetraploids in the 4*x*-6*x* populations (in Table 2) show specialization of dung saprotrophs (*Preussia spp*., *Podospora spp*., *Ascobolus spp. & Iodophanus spp*.) with low fidelity and perfect specificity. These fidelity and specificity values mean the OTUs are not associated with every tetraploid, but they are only present in soil associated with tetraploids. Dung saprotrophs make nutrients accessible that would otherwise be inaccessible to the host plant and can produce secondary metabolites that may act as antifungal agents against other fungi (Sarrocco 2016). For example, species of *Podospora* (Table 2) have been documented to produce at least seven different metabolites that have antifungal properties against genera that contain plant pathogens, *Fusarium* and *Ascobolus* (Sarrocco 2016). Antifungal functions gained from these OTUs’ presence may help tetraploid plants gain a fitness advantage due to a lower pathogen load. Another possible hypothesis is that tetraploid *L. tridentata* can access nutrients liberated by these dung saprotrophs that are not accessible to hexaploids, resulting in niche differences between the co-occurring cytotypes. The apparent replication of function (four different dung saprotrophs, from three different families found in association with tetraploids, but not hexaploids) is consistent with an area of niche space being partitioned based on specific interactions with the tetraploid hosts.

The indicator OTU combinations (Tables 3 & 4) are a source for further conjecture about MMND. All the indicator OTU combinations have moderate specificity and fidelity values, indicating they are present in most of the rhizosphere fungal assemblages of their respective cytotypes. Both mixed cytotype populations have indicator OTU combinations with a plant pathogen/wood saprotroph and a litter/soil saprotroph constituent. The repeated occurrence of these OTU combinations may indicate that they have an uncharacterized association or interdependency that may be important for the root economics spectrum of *L. tridentata* (Lugo, Anton, and Cabello 2005; Nguyen et al. 2011; Medeiros et al. 2021).

### Caveats

A major assumption of our study is that contemporary plant-fungal interactions are informative about past processes that have contributed to cytotype co-occurrence. Post-polyploidization evolutionary change and adaptation are likely to have occurred since the formation of tetraploid and hexaploid *L. tridentata* (Laport and Ramsey 2015; Walters and Freeman 1983), which may have also influenced cytotype-specific fungal associations. Thus, the fungal associate differences we documented here, and any MMND resulting from such differences, does not necessarily reflect the interspecific interactions that were important historically during the formation and establishment of the tetraploid and hexaploid cytotypes. Further, characterizing the population dynamics of long-lived perennials is challenging. The timeframe over which *L. tridentata* plants live makes it difficult to infer the fitness and niche divergence consequences of differences in fungal associates when typical population dynamics may unfold over centuries or millennia (Cody 2000).

In addition to uncertainty over whether contemporary fungal associate differences contributed to historical cytotype niche differentiation, it is also unclear whether seasonal rhizosphere dynamics may influence the population dynamics of *L. tridentata*. We collected rhizosphere samples at a single point in time, and thus these samples represent a narrow window into potentially complex rhizosphere fungal assemblage dynamics. The strong and varied seasonality of rainfall patterns in the Chihuahuan, Sonoran, and Mojave Deserts suggest the possibility of substantial fungal assemblage turnover over time (Clark, Rillig, and Nowak 2009). Relic DNA in soil has been shown to hinder the detection of temporal dynamics for soil microbial communities and clouds estimates of soil microbial diversity (Carini et al. 2017; 2020). Recent studies have documented seasonal turnover in rhizosphere fungal communities on diploid and tetraploid *Salicornia* (Gonçalves, Pena, and Albach 2022), which could play an important role in facilitating polyploid population dynamics.

### Future directions

Ploidy-specific soil microbiome differences may be important determinants of ecological performance and relative fitness in mixed-cytotype populations. Although we found that some of the rhizosphere OTUs differed between diploid, tetraploid, and hexaploid *L. tridentata*, we also found that the overall rhizosphere assemblages were similar. As with other studies of polyploid soil microbiomes, it remains unclear how important fungal associate differences might be in contributing to the MMND necessary to ease MCE and facilitate cytotype co-occurrence.

Manipulative experiments employing a plant-soil feedback design (Smith-Ramesh & Reynolds 2017) focused on co-occurring intra-specific cytotypes rather than co-occurring species have the potential to reveal the strength of rhizosphere-mediated MMND. Such experiments may also prove useful for predicting long-term population dynamics in polyploid species by revealing otherwise cryptic inter-specific ecological interactions that have only been accounted for indirectly in other studies of polyploid species. For example, accounting for microbially-mediated niche differences would help predict whether one cytotype is likely to overtake the other in the population or whether cytotype coexistence is likely.

Coupled with rhizosphere community sequencing, OTU functional group characterizations could also aid investigations into how whole genome duplication alters plant traits and patterns of biodiversity (K. A. Segraves 2017; Laport and Ng 2017). For example, our finding that soil fungal OTU identities differ strongly between the diploid-tetraploid mixed site and the tetraploid-hexaploid mixed site suggests landscape-level processes are important for determining soil fungal assemblages. Yet, it also appears that different OTUs may be involved in unique interactions among diploids and tetraploids than those observed for tetraploids and hexaploids.

These differences need additional study, such knowledge may also help us to understand the susceptibility of community-level biotic interactions and structure to increasing ecosystem disturbance and the persistence of native species in the face of non-native species introductions and climate change (K. A. Segraves and Anneberg 2016). What is clear is that soil fungal associate assemblages of polyploid species may be more complex and consequential than previously thought, and additional investigations of soil microbiomes and their interactions with polyploid species are needed to better quantify their effects on mediating community dynamics.

## Speculations

Another way to contextualize our study is to ask whether the fungal associations we document are consistent with *L. tridentata* cytotypes representing separate units of biodiversity. Biologists have long disagreed over whether the inter-cytotype reproductive isolation associated with genome duplication means cytotypes are different species, especially in the face of variable and often overlapping levels of phenotypic variation among cytotypes making their circumscription challenging. Unique associations with pollinators, herbivores, and soil microbes supports the opinion that widespread ploidal races represent functional units of biodiversity. As in similar studies, *L. tridentata* cytotypes vary across multiple ecological axes and exhibit the distinct community-level and inter-specific interactions observed among traditionally circumscribed taxa. Such differences are informative about community structure and the functioning of ecological systems. Differential associations between plants differing in ploidy and rhizosphere OTUs represents variation along multiple axes of a Hutchinsonian *n*-dimensional niche hypervolume, and perhaps should be considered no less important from a biodiversity perspective than the partitioning of foraging spaces in spruce canopies by some warblers in the northeastern U.S. Recognition of ploidal diversity, and the diversity of traits exhibited by different cytotypes, would appropriately respect the biodiversity implications of a ubiquitous evolutionary process and its ecological consequences.

## Data Archiving Statement

Data, metadata and associated analysis scripts will be deposited with DRYAD, prior to final publication.

## Ethics Statement

We acknowledge field collections were conducted on lands that are the traditional territories of the Cocopah, O’odham Jewed, Sobaipuri, Ndee and Hohokam people. The authors reside and work on lands that are the traditional territories of the Tiwa, Piro, Puebloan, Numu, and Shoshone-Bannock. We wish to express our gratitude and respect for these people and their stewardship of the land. Forced dispossession and deliberate colonization through direct coercion, treaty and war has been United States governmental practice for centuries. We as scientists and our institutions continue to benefit from these practices. To ignore that is to perpetuate injustice to populations of people that no longer exist on these lands. We understand that territorial acknowledgement is only a gesture, but it represents the beginning of our commitment to justice and reconciliation in the United States.

## Acknowledgements

We would like to thank Purbendra Yogi for his tireless support and assistance in the lab. BG thanks their dissertation committee members Drs. Helen Wearing and Jenn Rudgers for feedback that strengthened the project design.

---

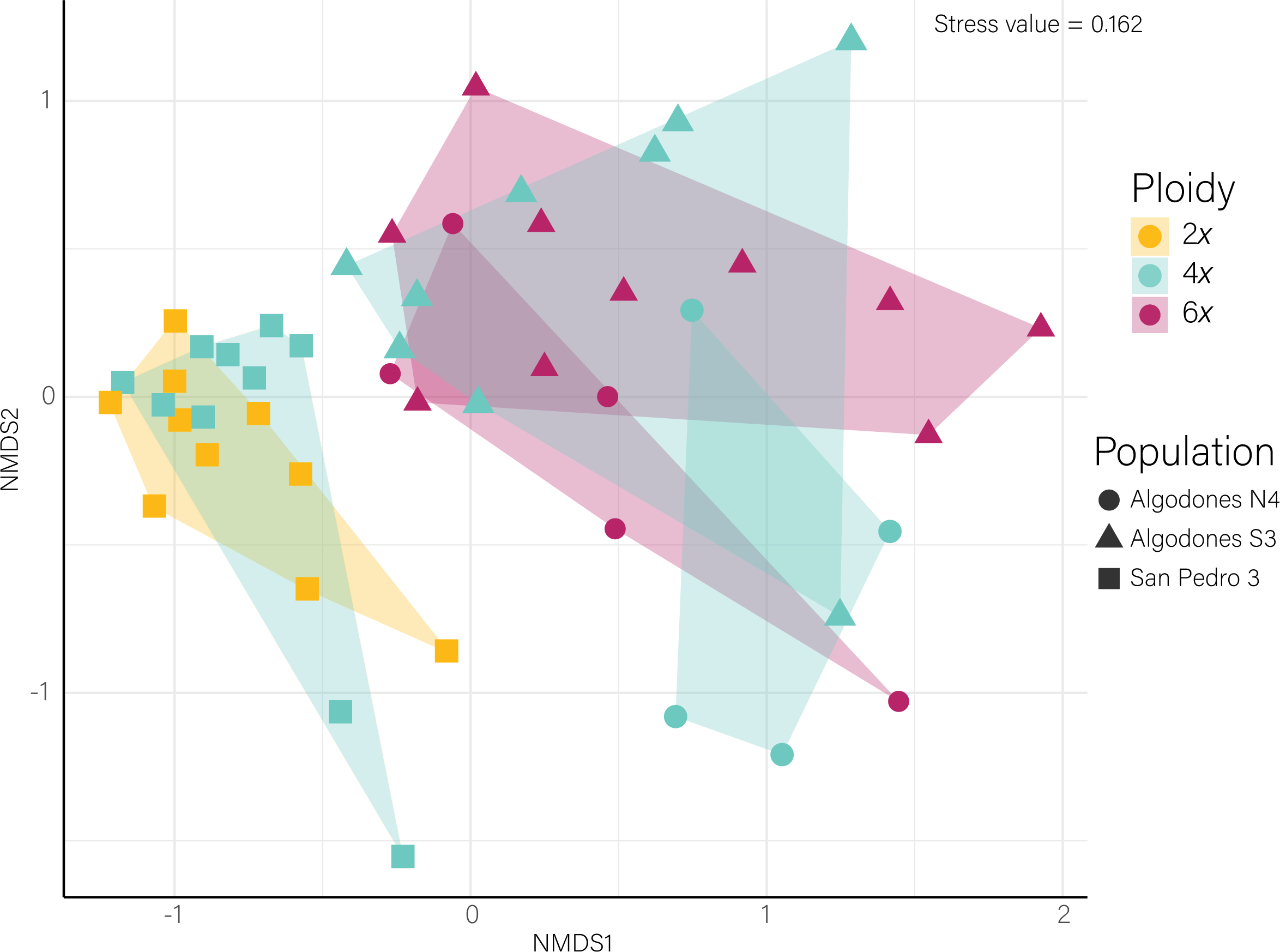

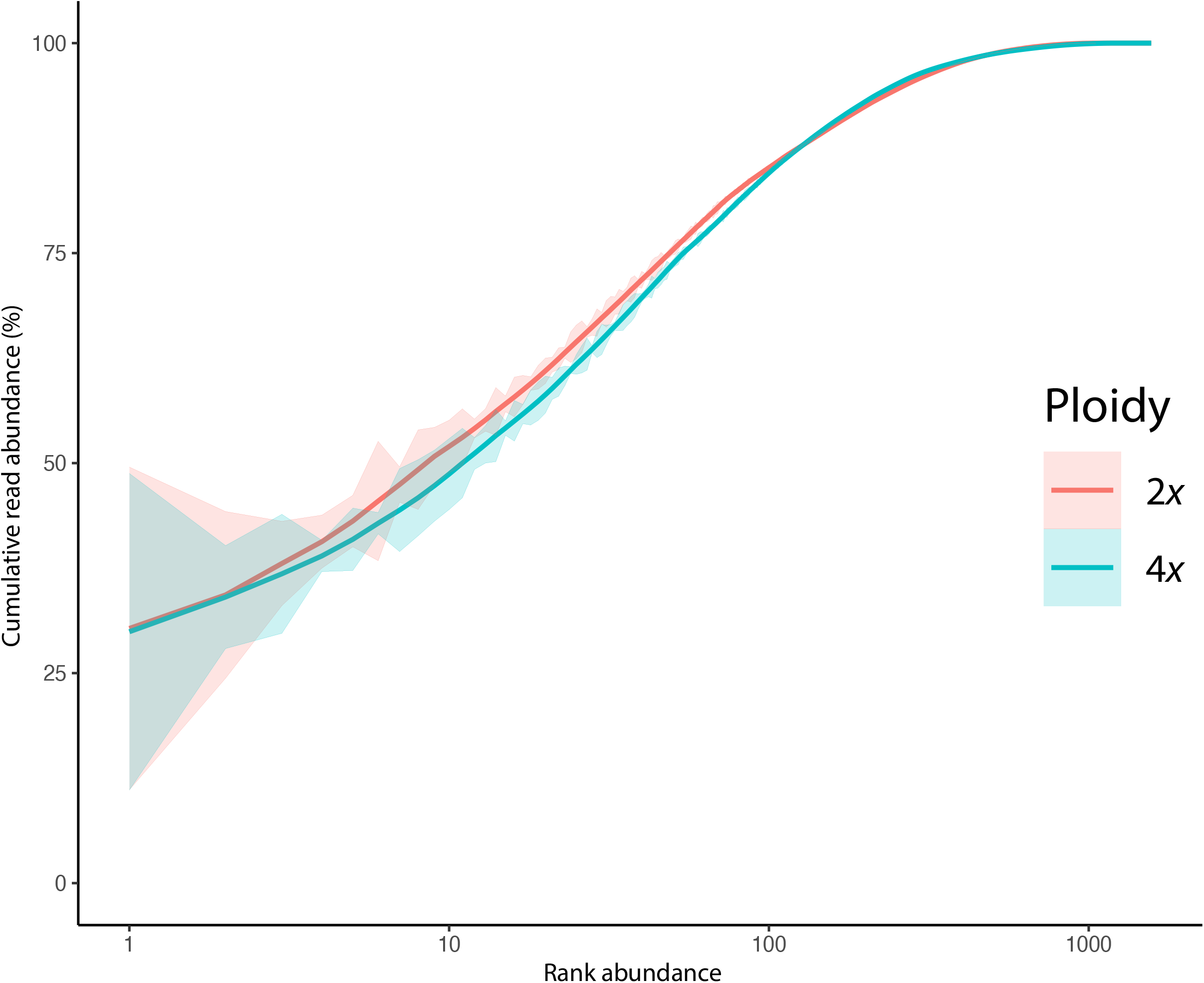

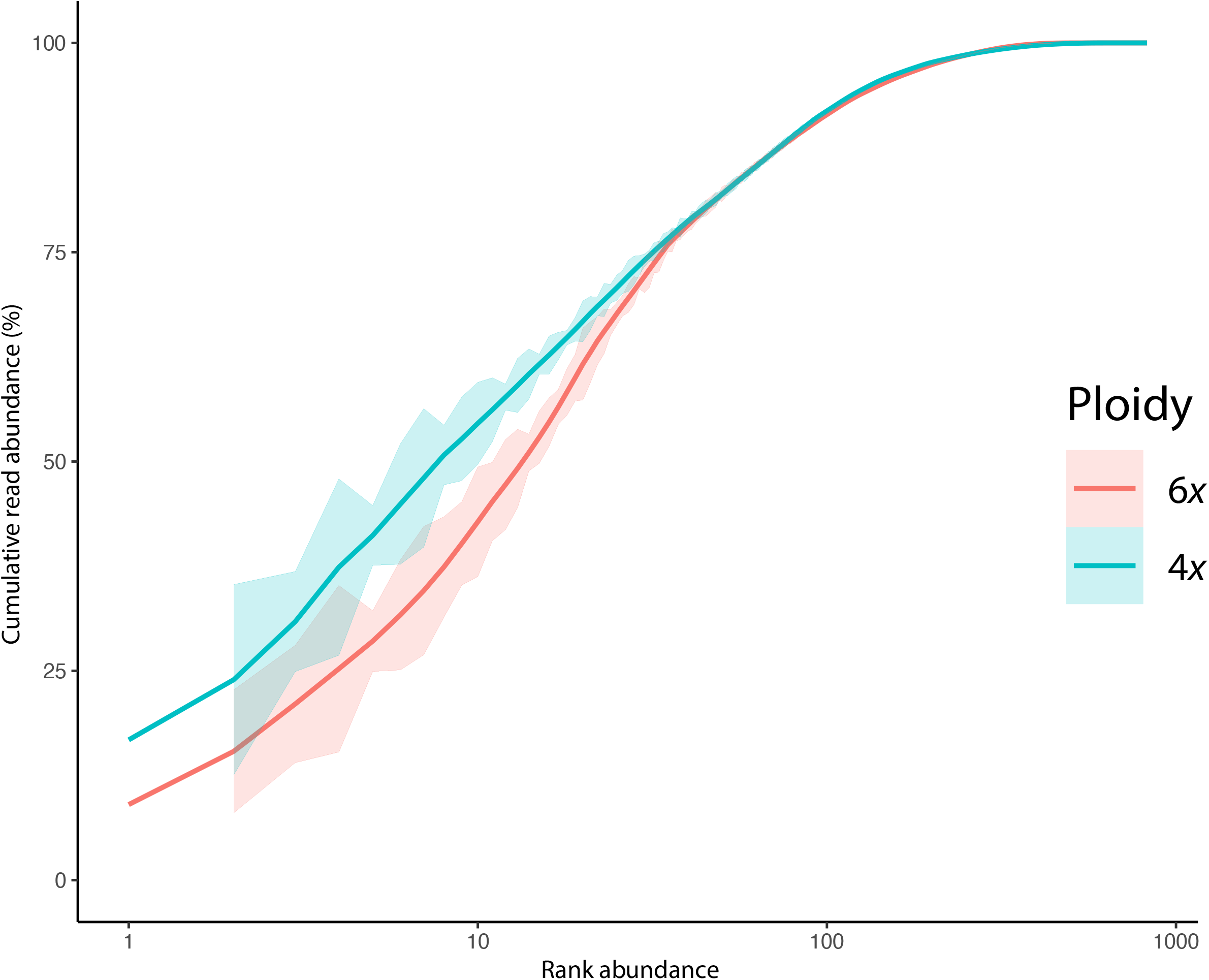

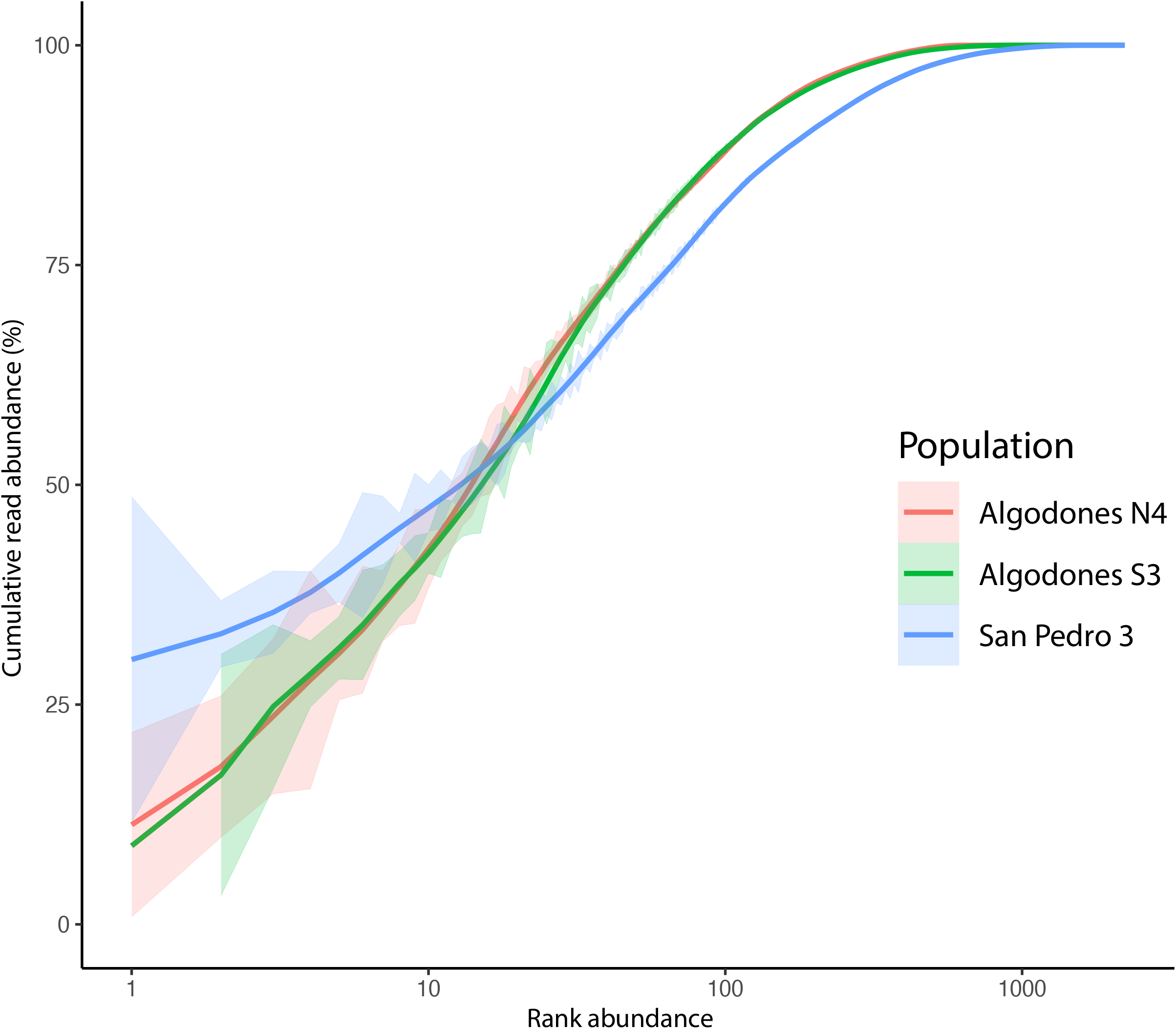

## Bibliography

Abarenkov, Kessy, Allan Zirk, Timo Piirmann, Raivo Pöhönen, Filipp Ivanov, R. Henrik Nilsson, and Urmas Kõljalg. 2021. “UNITE General FASTA Release for Eukaryotes 2.” R package. UNITE Community. https://doi.org/10.15156/BIO/1280160.

Andersen, K.S., R.H. Kirkgaard, S.M. Karst, and M. Albertsen. 2018. “Ampvis2: An R Package to Analyse and Visualise 16S RRNA Amplicon Data.” R. https://doi.org/10.1101/299537.

Bouffaud, M.-L., C. Bragalini, A. Berruti, M. Peyret-Guzzon, S. Voyron, H. Stockinger, D. van Tuinen, et al. 2017. “Arbuscular Mycorrhizal Fungal Community Differences among European Long-Term Observatories.” Mycorrhiza 27 (4): 331–43. https://doi.org/10.1007/s00572-016-0753-9.

Cáceres Miquel De, and Pierre Legendre. 2009. “Associations between Species and Groups of Sites: Indices and Statistical Inference.” Ecology 90 (12): 3566–74. https://doi.org/10.1890/08-1823.1.

Carini, Paul, Manuel Delgado-Baquerizo, Eve-Lyn S. Hinckley, Hannah Holland-Moritz, Tess E. Brewer, Garrett Rue, Caihong Vanderburgh, Diane McKnight, and Noah Fierer. 2020. “Effects of Spatial Variability and Relic DNA Removal on the Detection of Temporal Dynamics in Soil Microbial Communities.” Edited by Janet K. Jansson. MBio 11 (1): e02776–19. https://doi.org/10.1128/mBio.02776-19.

Carini, Paul, Patrick J. Marsden, Jonathan W. Leff, Emily E. Morgan, Michael S. Strickland, and Noah Fierer. 2017. “Relic DNA Is Abundant in Soil and Obscures Estimates of Soil Microbial Diversity.” Nature Microbiology 2 (3): 16242. https://doi.org/10.1038/nmicrobiol.2016.242.

Čertner, Martin, Pavel Kúr, Filip Kolář, and Jan Suda. 2019. “Climatic Conditions and Human Activities Shape Diploid–Tetraploid Coexistence at Different Spatial Scales in the Common Weed Tripleurospermum Inodorum (Asteraceae).” Journal of Biogeography 46 (7): 1355–66. https://doi.org/10.1111/jbi.13629.

Clark, N.M., M.C. Rillig, and R.S. Nowak. 2009. “Arbuscular Mycorrhizal Fungal Abundance in the Mojave Desert: Seasonal Dynamics and Impacts of Elevated CO2.” Journal of Arid Environments 73 (9): 834–43. https://doi.org/10.1016/j.jaridenv.2009.03.004.

Cody, Martin L. 2000. “Slow-motion Population Dynamics in Mojave Desert Perennial Plants.” Journal of Vegetation Science 11 (3): 351–58. https://doi.org/10.2307/3236627.

Coyne, Jerry, and Alan Orr. 2004. Speciation. Sinauer Associates.

De Cáceres, Miquel, Pierre Legendre, Susan K. Wiser, and Lluís Brotons. 2012. “Using Species Combinations in Indicator Value Analyses.” Edited by Robert B. O’Hara. Methods in Ecology and Evolution 3 (6): 973–82. https://doi.org/10.1111/j.2041-210X.2012.00246.x.

Decanter, Lucile, Guy Colling, Nora Elvinger, Starri Heiðmarsson, and Diethart Matthies. 2020. “Ecological Niche Differences between Two Polyploid Cytotypes of Saxifraga Rosacea.” American Journal of Botany 107 (3): 423–35. https://doi.org/10.1002/ajb2.1431.

Dufrêne, Marc, and Pierre Legendre. 1997. “Species Assemblages and Indicatopr Species: The Need for a Flexible Aysmmetrical Approach.” Ecological Monographs 67 (3): 345–66. https://doi.org/10.1890/0012-9615(1997)067[0345:SAAIST]2.0.CO;2.

Edgar, E.C. 2013. “UPARSE: Highly Accurate OTU Sequences from Microbial Amplicon Reads.” Nature Methods. https://www.nature.com/articles/nmeth.2604.

Edgar, Robert C. 2010. “Search and Clustering Orders of Magnitude Faster than BLAST.” Bioinformatics 26 (19): 2460–61. https://doi.org/10.1093/bioinformatics/btq461.

Edgar, Robert C.. 2016. “SINTAX: A Simple Non-Bayesian Taxonomy Classifier for 16S and ITS Sequences.” Preprint. Bioinformatics. https://doi.org/10.1101/074161.

Fowler, Norma L., and Donald A. Levin. 2016. “Critical Factors in the Establishment of Allopolyploids.” American Journal of Botany 103 (7): 1236–51. https://doi.org/10.3732/ajb.1500407.

Gaynor, Michelle L., D. Blaine Marchant, Douglas E. Soltis, and Pamela S. Soltis. 2018. “Climatic Niche Comparison among Ploidal Levels in the Classic Autopolyploid System, Galax Urceolata.” American Journal of Botany 105 (10): 1631–42. https://doi.org/10.1002/ajb2.1161.

Gonçalves, Danilo Reis, Rodica Pena, and Dirk C. Albach. 2022. “Polyploidy and Plant-Fungus Symbiosis: Evidence of Cytotype-Specific Microbiomes in the Halophyte Salicornia (Amaranthaceae).” Preprint. bioRxiv. https://doi.org/10.1101/2022.03.09.483717.

Hunter, Kimberly L., Julio L. Betancourt, Brett R. Riddle, Thomas R. Van Devender, Kenneth L. Cole, and W. Geoffrey Spaulding. 2001. “Ploidy Race Distributions since the Last Glacial Maximum in the North American Desert Shrub, Larrea Tridentata: Ploidy Race Distributions in Larrea Tridentata.” Global Ecology and Biogeography 10 (5): 521–33. https://doi.org/10.1046/j.1466-822X.2001.00254.x.

Husband, Brian. 2000. “Constraints on Polyploid Evolution: A Test of the Minority Cytotype Exclusion Principle,” 7.

Husband, Brian C., Barbara Ozimec, Sara L. Martin, and Lisa Pollock. 2008. “Mating Consequences of Polyploid Evolution in Flowering Plants: Current Trends and Insights from Synthetic Polyploids.” International Journal of Plant Sciences 169 (1): 195–206. https://doi.org/10.1086/523367.

Landis, Jacob B., Douglas E. Soltis, Zheng Li, Hannah E. Marx, Michael S. Barker, David C. Tank, and Pamela S. Soltis. 2018. “Impact of Whole-Genome Duplication Events on Diversification Rates in Angiosperms.” American Journal of Botany 105 (3): 348–63. https://doi.org/10.1002/ajb2.1060.

Laport, Robert G., Layla Hatem, Robert L. Minckley, and Justin Ramsey. 2013. “Ecological Niche Modeling Implicates Climatic Adaptation, Competitive Exclusion, and Niche Conservatism among Larrea Tridentata Cytotypes in North American Deserts 1, 2.” The Journal of the Torrey Botanical Society 140 (3): 349–63. https://doi.org/10.3159/TORREY-D-13-00009.1.

Laport, Robert G., Robert L. Minckley, and Diana Pilson. 2021. “Pollinator Assemblage and Pollen Load Differences on Sympatric Diploid and Tetraploid Cytotypes of the Desert-dominant Larrea Tridentata.” American Journal of Botany 108 (2): 297–308. https://doi.org/10.1002/ajb2.1605.

Laport, Robert G., Robert L. Minckley, and Justin Ramsey. 2012. “Phylogeny and Cytogeography of the North American Creosote Bush (<I>Larrea Tridentata</I>, Zygophyllaceae).” Systematic Botany 37 (1): 153–64. https://doi.org/10.1600/036364412X616738.

Laport, Robert G., Robert L. Minckley, and Justin Ramsey. 2016. “Ecological Distributions, Phenological Isolation, and Genetic Structure in Sympatric and Parapatric Populations of the Larrea Tridentata Polyploid Complex.” American Journal of Botany 103 (7): 1358–74. https://doi.org/10.3732/ajb.1600105.

Laport, Robert G., and Julienne Ng. 2017. “Out of One, Many: The Biodiversity Considerations of Polyploidy.” American Journal of Botany 104 (8): 1119–21. https://doi.org/10.3732/ajb.1700190.

Laport, Robert G., and Justin Ramsey. 2015. “Morphometric Analysis of the North American Creosote Bush (Larrea Tridentata, Zygophyllaceae) and the Microspatial Distribution of Its Chromosome Races.” Plant Systematics and Evolution 301 (6): 1581–99. https://doi.org/10.1007/s00606-014-1179-5.

Levin, Donald A. 1975. “Minority Cytotype Exclusion in Local Plant Populations.” Taxon 24 (1): 35–43.

Levin, Donald A. 1983. “Polyploidy and Novelty in Flowering Plants.” The American Naturalist 122 (1): 1–25. https://doi.org/10.1086/284115.

López-Jurado, Javier, Enrique Mateos-Naranjo, and Francisco Balao. 2019. “Niche Divergence and Limits to Expansion in the High Polyploid Dianthus Broteri Complex.” New Phytologist 222 (2): 1076–87. https://doi.org/10.1111/nph.15663.

Lugo, M A, A M Anton, and M N Cabello. 2005. “Arbuscular Mycorrhizas in the Larrea Divaricata Scrubland of the Arid ‘Chaco’, Central Argentina,” 16.

Mabry, T.J., J.H. Hunziker, and D.R. Difeo, Jr. 1977. Creosote Bush: Biology and Chemistry of Larrea in New World Deserts. 1st ed. US/IBP Synthesis Series 6. John Wiley & Sons, Inc.

MacArthur, Robert. 1958. “Population Ecology of Some Warblers of Northeastern Coniferous Forests.” Ecology 39 (4): 599–619.

Maherali, Hafiz, Alison Walden, and Brian Husband. 2009. “Genome Duplication and the Evolution of Physiological Responses to Water Stress.” New Phytologist 184: 721–31.

Martin, Marcel. 2011. “Cutadapt Removes Adapter Sequences from High-Throughput Sequencing Reads.” EMB.Net Journal 17 (May): 3. https://doi.org/10.14806/ej.17.1.200.

Medeiros, Lucas P., Karina Boege, Ek del-Val, Alejandro Zaldívar-Riverón, and Serguei Saavedra. 2021. “Observed Ecological Communities Are Formed by Species Combinations That Are among the Most Likely to Persist under Changing Environments.” The American Naturalist 197 (1): E17–29. https://doi.org/10.1086/711663.

Muñoz-Pajares, A. J., F. Perfectti, J. Loureiro, M. Abdelaziz, P. Biella, M. Castro, S. Castro, and J. M. Gómez. 2018. “Niche Differences May Explain the Geographic Distribution of Cytotypes in Erysimum Mediohispanicum.” Plant Biology 20 (Stebbins 1971): 139–47. https://doi.org/10.1111/plb.12605.

Münzbergová, Zuzana, Jiří Skuhrovec, and Petr Maršík. 2015. “Large Differences in the Composition of Herbivore Communities and Seed Damage in Diploid and Autotetraploid Plant Species.” Biological Journal of the Linnean Society 115 (2): 270–87. https://doi.org/10.1111/bij.12482.

Nguyen, L.M., M.P. Buttner, P. Cruz, S.D. Smith, and E.A. Robleto. 2011. “Effects of Elevated Atmospheric CO2 on Rhizosphere Soil Microbial Communities in a Mojave Desert Ecosystem.” Journal of Arid Environments 75 (10): 917–25. https://doi.org/10.1016/j.jaridenv.2011.04.028.

O’Connor, Timothy K., Robert G. Laport, and Noah K. Whiteman. 2019. “Polyploidy in Creosote Bush (Larrea Tridentata) Shapes the Biogeography of Specialist Herbivores.” Journal of Biogeography 46 (3): 597–610. https://doi.org/10.1111/jbi.13490.

Oksanen, Jari, F. Guillaume Blanchet, Michael Friendly, Roeland Kindt, Pierre Legendre, Dan McGlinn, Peter R. Minchin, et al. 2020. “Vegan.” R package. https://CRAN.R-project.org/package=vegan.

Põlme, Sergei, Kessy Abarenkov, R. Henrik Nilsson, Björn D. Lindahl, Karina Engelbrecht Clemmensen, Havard Kauserud, Nhu Nguyen, et al. 2020. “FungalTraits: A User-Friendly Traits Database of Fungi and Fungus-like Stramenopiles.” Fungal Diversity 105 (1): 1–16. https://doi.org/10.1007/s13225-020-00466-2.

R Core Team. 2022. “R: A Language and Evironment for Statistical Computing.” R. Vienna, Austria: R Foundation for Statistical Computing. https://www.R-project.org/.

Ramsey, Justin, and Tara S. Ramsey. 2014. “Ecological Studies of Polyploidy in the 100 Years Following Its Discovery.” Philosophical Transactions of the Royal Society B: Biological Sciences 369 (1648): 20130352. https://doi.org/10.1098/rstb.2013.0352.

Salazar, Guillem. 2022. “EcoUtils.” R package. https://github.com/GuillemSalazar/EcolUtils.

Sarrocco, Sabrina. 2016. “Dung-Inhabiting Fungi: A Potential Reservoir of Novel Secondary Metabolites for the Control of Plant Pathogens: Antifungal Secondary Metabolites from Coprophilous Fungi.” Pest Management Science 72 (4): 643–52. https://doi.org/10.1002/ps.4206.

Segraves, Kari A. 2017. “The Effects of Genome Duplications in a Community Context.” New Phytologist 215 (1): 57–69. https://doi.org/10.1111/nph.14564.

Segraves, Kari A., and Thomas J. Anneberg. 2016. “Species Interactions and Plant Polyploidy.” American Journal of Botany 103 (7): 1326–35. https://doi.org/10.3732/ajb.1500529.

Segraves, Kari, and J. Thompson. 1999. “Plant Polyploidy and Pollination: Floral Traits and Insect Visits to Diploid and Tetraploid Heuchera Grossulariifolia.” Evolution 53 (4): 1114–27.

Sudova, R., H. Pankova, J. Rydlova, Z. Munzbergova, and J. Suda. 2014. “Intraspecific Ploidy Variation: A Hidden, Minor Player in Plant-Soil-Mycorrhizal Fungi Interactions.” American Journal of Botany 101 (1): 26–33. https://doi.org/10.3732/ajb.1300262.

Sudová, Radka, Petr Kohout, Zuzana Kolaříková, Jana Rydlová, Jana Voříšková, Jan Suda, Stanislav Španiel, Heinz Müller-Schärer, and Patrik Mráz. 2018. “Sympatric Diploid and Tetraploid Cytotypes of Centaurea Stoebe s.l. Do Not Differ in Arbuscular Mycorrhizal Communities and Mycorrhizal Growth Response.” American Journal of Botany, December, ajb2.1206. https://doi.org/10.1002/ajb2.1206.

Taylor, D. Lee, William A. Walters, Niall J. Lennon, James Bochicchio, Andrew Krohn, J. Gregory Caporaso, and Taina Pennanen. 2016. “Accurate Estimation of Fungal Diversity and Abundance through Improved Lineage-Specific Primers Optimized for Illumina Amplicon Sequencing.” Edited by D. Cullen. Applied and Environmental Microbiology 82 (24): 7217–26. https://doi.org/10.1128/AEM.02576-16.

Těšitelová, Tamara, Jana Jersáková, Mélanie Roy, Barbora Kubátová, Jakub Těšitel, Tomáš Urfus, Pavel TrávníČek, and Jan Suda. 2013. “Ploidy-Specific Symbiotic Interactions: Divergence of Mycorrhizal Fungi between Cytotypes of the Gymnadenia Conopsea Group (Orchidaceae).” New Phytologist 199 (4): 1022–33. https://doi.org/10.1111/nph.12348.

Walters, James Peter, and C. Edward Freeman. 1983. “Growth Rates and Root: Shoot Ratios in Seedlings of the Desert Shrub Larrea Tridentata.” The Southwestern Naturalist 28 (3): 357. https://doi.org/10.2307/3670798.

Wan, J. Z., L. X. Chen, S. Gao, Y. B. Song, S. L. Tang, F. H. Yu, J. M. Li, and M. Dong. 2019. “Ecological Niche Shift between Diploid and Tetraploid Plants of Fragaria (Rosaceae) in China.” South African Journal of Botany 121: 68–75. https://doi.org/10.1016/j.sajb.2018.10.027.

Wan, J.-Z., L.-X. Chen, S. Gao, Y.-B. Song, S.-L. Tang, F.-H. Yu, J.-M. Li, and M. Dong. 2019. “Ecological Niche Shift between Diploid and Tetraploid Plants of Fragaria (Rosaceae) in China.” South African Journal of Botany 121 (March): 68–75. https://doi.org/10.1016/j.sajb.2018.10.027.

Wickham, H. 2016. “Ggplot2.” R package. Springer-Verlag.

Wickham, Hadley, Mara Averick, Jennifer Bryan, Winston Chang, Lucy D’ Agostino McGowan, Romain François, Garrett Grolemund, et al. 2019. “Welcome to the Tidyverse.” R package. https://doi.org/10.21105/joss.01686.

Wood, T. E., N. Takebayashi, M. S. Barker, I. Mayrose, P. B. Greenspoon, and L. H. Rieseberg. 2009. “The Frequency of Polyploid Speciation in Vascular Plants.” Proceedings of the National Academy of Sciences 106 (33): 13875–79. https://doi.org/10.1073/pnas.0811575106.

Wu, Shuqi, Jiliang Cheng, Xinyu Xu, Yi Zhang, Yexin Zhao, Huixin Li, and Sheng Qiang. 2019. “Polyploidy in Invasive Solidago Canadensis Increased Plant Nitrogen Uptake, and Abundance and Activity of Microbes and Nematodes in Soil.” Soil Biology and Biochemistry 138 (November): 107594. https://doi.org/10.1016/j.soilbio.2019.107594.

