## Supplemental Table and Figures for "Differentiation of rhizosphere fungal assemblages by host ploidy level in mixed-ploidy *Larrea tridentata* populations"

Supplement:

Table S1: PERMANOVA test results (p-values) for different models for rhizosphere assemblage composition explained by Ploidy, Population and Ploidy*Population. Filtering value is the threshold used for filtering total read abundance for the OTU to be included. Across all models population is the only significant predictor, supporting the idea that rhizosphere fungal assemblages are more similar at the population level than the same ploidies are in different populations. Significant p-values are italicized.

| **Type of Error Calculation** | | | **Marginal Effects** | **Sequential** | | | |
|---|---|---|---|---|---|---|---|
| **Filtering Value** | **Number of OTU's** | **Model Terms** | **Full Model** ~ Ploidy+Pop + Ploidy*Pop | **Full Model** ~ Ploidy+Pop + Ploidy*Pop | **Full Model** ~ Pop+Ploidy + Pop*Ploidy | **Reduced Model** ~ Ploidy+Pop | **Reduced Model** ~ Pop+Ploidy |
| 3800 | 2217 | **Ploidy** | - | 0.5668 | 0.5795 | 0.5877 | 0.5937 |
| | | **Population** | - | *0.00009* | *0.00009* | *0.00009* | *0.00009* |
| | | **Ploidy*Pop** | 0.1727 | 0.1714 | 0.1722 | - | - |
| 4000 | 2217 | **Ploidy** | - | 0.7050 | 0.6806 | 0.7135 | 0.698 |
| | | **Population** | - | *0.00009* | *0.00009* | *0.00009* | *0.00009* |
| | | **Ploidy*Pop** | 0.1592 | 0.1607 | 0.1644 | - | - |
| 12000 | 2172 | **Ploidy** | - | 0.4590 | 0.3928 | 0.469 | 0.4195 |
| | | **Population** | - | *0.00009* | *0.00009* | *0.00009* | *0.00009* |
| | | **Ploidy*Pop** | 0.1033 | 0.1039 | 0.1098 | - | - |
| 20000 | 2172 | **Ploidy** | - | 0.4486 | 0.3940 | 0.4684 | 0.4216 |
| | | **Population** | - | *0.00009* | *0.00009* | *0.00009* | *0.00009* |
| | | **Ploidy*Pop** | 0.109 | 0.1110 | 0.1076 | - | - |

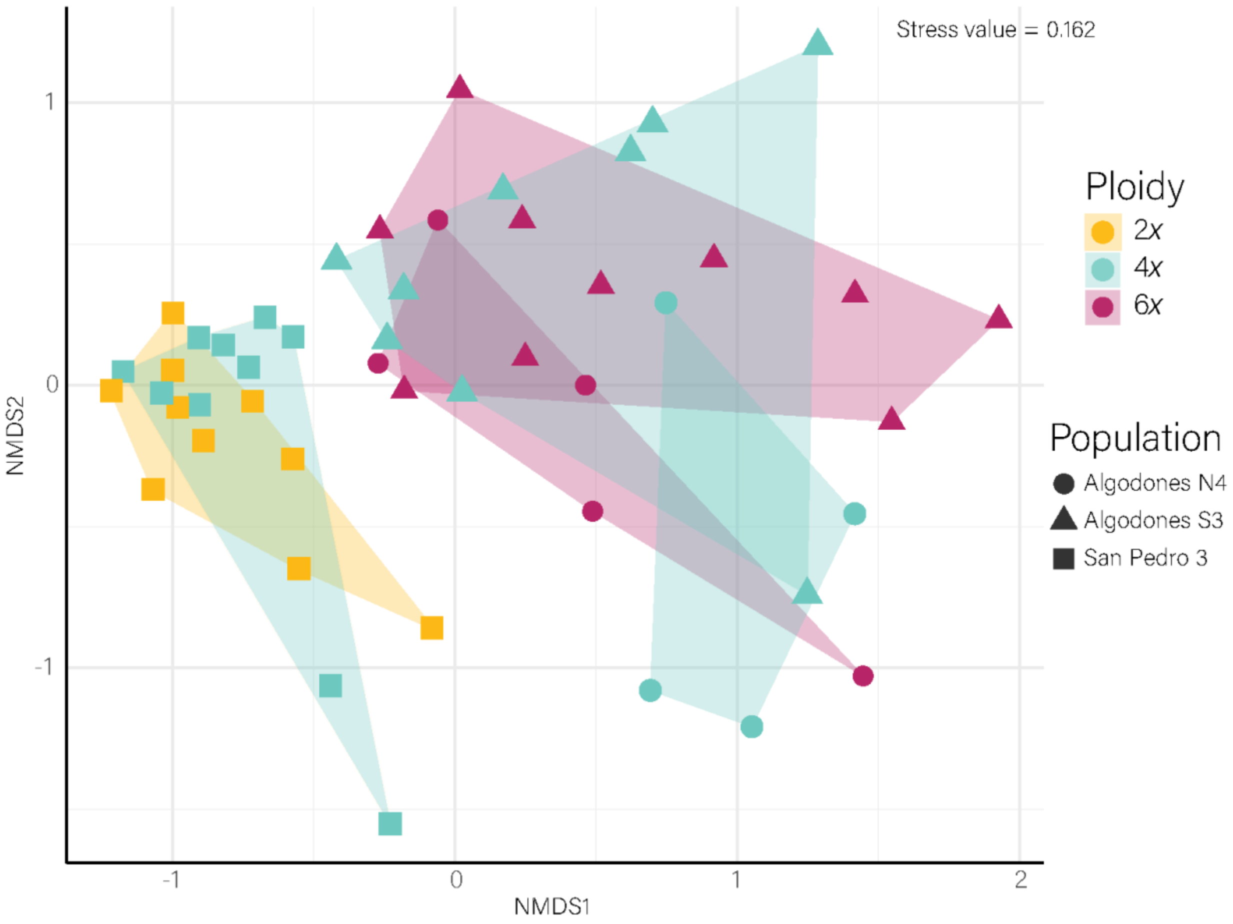


Figure S1: NMDS ordination for rhizosphere fungal assemblage composition of *Larrea tridentata* cytotypes in mixed cytotype populations. There is minimal overlap in rhizosphere fungal assemblage composition between 4*x* at Algodones N4 and Algodones S3. Otherwise cytotypes with the same population identity have the greatest overlap in rhizosphere fungal assemblage composition.

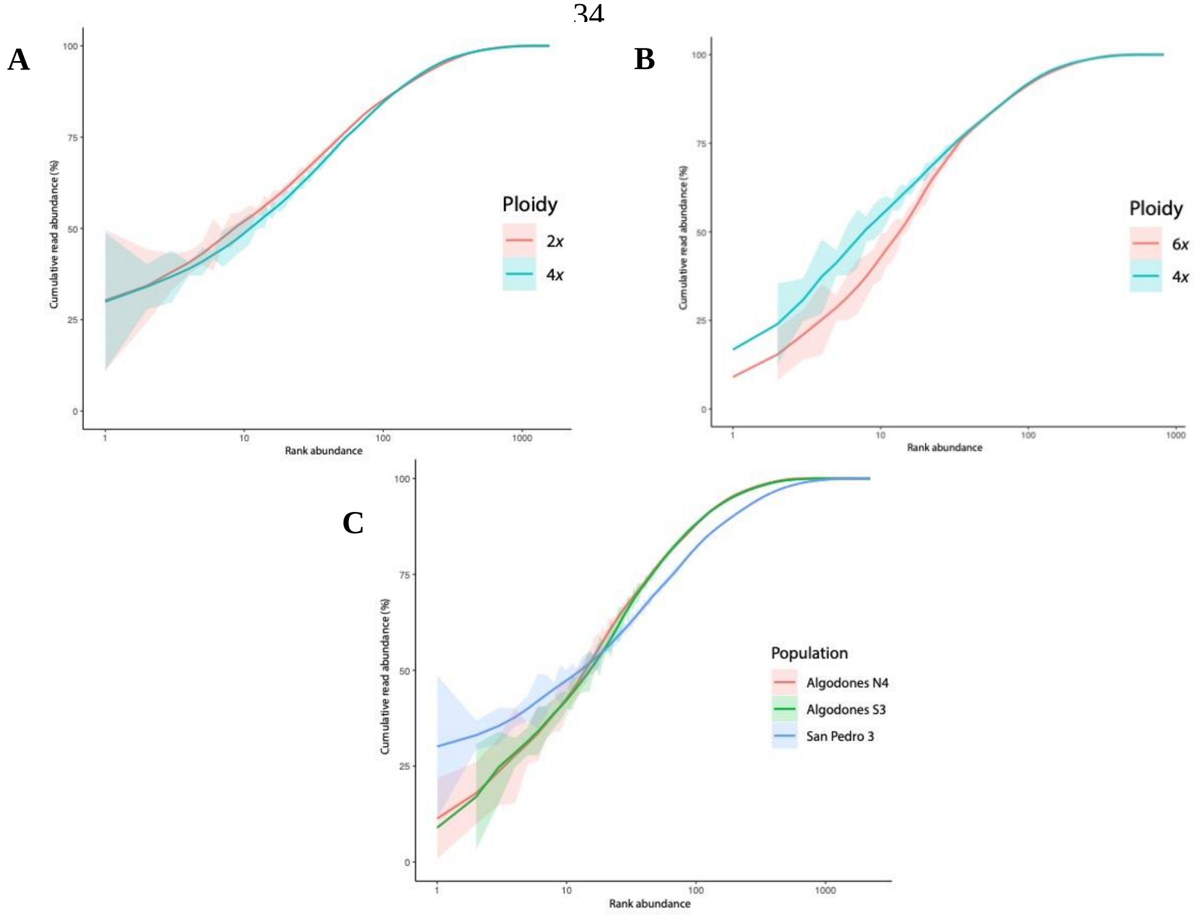


Figure S2: Cumulative read abundance by OTU rank abundance for **(A)** 2*x*-4*x*, **(B)** 4*x*-6*x* and **(C)** Population Identity. **(A)** shows read abundance does not differ between 2*x* and 4*x* cytotypes in the 2*x*-4*x* population (San Pedro 3). **(B)** shows 4*x* had a faster climb in read abundance for lower rank abundance values, meaning 4*x* had fewer OTUs making up their total reads (i.e., lower richness) in the 4*x*-6*x* populations. **(C)** shows read abundance differs between the San Pedro 3 population (2*x*-4*x*) and the Algodones N4 and Algodones S3 populations (4*x*-6*x*), which had approximately the same change in read abundance, meaning that the rhizospheres of the 2*x*-4*x* population had fewer OTUs making up their total reads (i.e., lower richness) than 4*x*-6*x* populations.
